# Gestational exposure to unmethylated CpG oligonucleotides dysregulates placental molecular clock network and fetoplacental growth dynamics, and disrupts maternal blood pressure circadian rhythms in rats

**DOI:** 10.1101/2023.03.14.532649

**Authors:** Jessica L. Bradshaw, Spencer C. Cushen, Contessa A. Ricci, Selina M. Tucker, Jennifer J. Gardner, Joel T. Little, Oluwatobiloba Osikoya, Styliani Goulopoulou

## Abstract

Bacterial infections and impaired mitochondrial DNA dynamics are associated with adverse pregnancy outcomes. Unmethylated cytosine-guanine dinucleotide (CpG) motifs are common in bacterial and mitochondrial DNA and act as potent immunostimulators. Here, we tested the hypothesis that exposure to CpG oligonucleotides (ODN) during pregnancy would disrupt blood pressure circadian rhythms and the placental molecular clock machinery, mediating aberrant fetoplacental growth dynamics. Rats were repeatedly treated with CpG ODN in the 3^rd^ trimester (gestational day, GD, 14, 16, 18) and euthanized on GD20 (near term) or with a single dose of CpG ODN and euthanized 4 hours after treatment on GD14. Hemodynamic circadian rhythms were analyzed via Lomb-Scargle periodogram analysis on 24-h raw data collected continuously via radiotelemetry. A *p*-value ≥ 0.05 indicates the absence of a circadian rhythm. Following the first treatment with CpG ODN, maternal systolic and diastolic blood pressure circadian rhythms were lost (*p* ≥ 0.05). Blood pressure circadian rhythm was restored by GD16 and remained unaffected after the second treatment with CpG ODN (*p* < 0.0001). Diastolic blood pressure circadian rhythm was again lost after the last treatment on GD18 (*p* ≥ 0.05). CpG ODN increased placental expression of *Per2* and *Per3* and *Tnfα* (*p* ≤ 0.05) and affected fetoplacental growth dynamics, such as reduced fetal and placental weights were disproportionately associated with increases in the number of resorptions in ODN-treated dams compared to controls. In conclusion, gestational exposure to unmethylated CpG DNA dysregulates placental molecular clock network and fetoplacental growth dynamics and disrupts blood pressure circadian rhythms.

## INTRODUCTION

During pregnancy, the maternal immune system undergoes adaptations that ensure tolerance to the fetal allograft while mothers maintain their ability to respond to exogenous pathogens (1). It is postulated that the interaction between endocrine and immune system adaptations during pregnancy often guides an aberrant maternal response to exogenous pathogens (2). As a result, pregnant women show greater susceptibility and worse clinical outcomes in response to bacterial and viral infections, such as *Listeria monocytogenes* and COVID19, compared to age-matched non-pregnant women (2–5). Moreover, infections and dysregulation of the immune system during pregnancy are associated with adverse cardio-obstetric outcomes such as gestational hypertension and preeclampsia, as well as other obstetric complications including preterm birth, intrauterine growth restriction, and stillbirth (2, 6–9).

Unmethylated cytosine-phosphodiester-guanine (CpG) motifs are potent triggers of host immune responses via stimulation of toll-like receptor 9 (TLR9), which is a pattern recognition receptor of the innate immune system (10–13). These unmethylated motifs are enriched in microbial DNA, mitochondrial DNA (mtDNA), and fetal DNA; yet these modified sequences are rare in mammalian nuclear DNA (10). Additionally, synthetic CpG oligonucleotides (ODN) are increasingly used in pre-clinical studies and ongoing clinical trials as effective vaccine adjuvants for mucosal vaccines against infectious diseases and allergens (14–20). However, these studies and trials are devoid in examining the effects of CpG ODN on maternal outcomes during pregnancy. Notably, concentrations of cell-free mtDNA in maternal sera increase during healthy human pregnancy and return to non-pregnant state values within six-eight weeks postpartum (21). Thus, elevations in mtDNA and presence of fetal DNA during pregnancy increase endogenous sources of unmethylated CpG sequences in pregnant women and may modulate immune responses and contribute to perinatal outcomes. In addition to microbial infections (i.e., exogenous immune triggers), circulating cell-free mtDNA and fetal DNA (i.e., endogenous damage-associated molecular patterns) have been implicated in obstetric complications such as preeclampsia and preterm birth (11, 22–24).

Cardiovascular functions such as blood pressure and heart rate follow a circadian pattern, exhibiting an increase during the wake phase (rodents), or mornings (humans), and a decrease during the sleep phase, or nights, respectively (25). Previous studies have shown that circadian patterns of blood pressure are stronger predictors of cardiovascular events compared to averaged daily blood pressure measurements in young patients with hypertension (26). Importantly, disruption of the circadian rhythm during pregnancy adversely affects fetal development (27) and is associated with pregnancy complications, such as intrauterine growth restriction, preeclampsia, HELLP (**h**emolysis, **e**levated **l**iver enzymes, and **l**ow **p**latelets) syndrome, and preterm birth, which have long-lasting effects on maternal and offspring health outcomes (28–30). Others have established an association between blood pressure circadian patterns and markers of immune system activation in non-pregnant patients with autoimmune disorders (31).

Circadian rhythms are controlled by “clock” genes, which are regulated by mediators of proinflammatory innate immune system responses (32–34). Clock genes have been identified in the placenta, and circadian disruption is associated with compromised placental function and fetal growth restriction (35, 36). Still, an association between immune system dysregulation during pregnancy, which is a feature of many pregnancy complications, and circadian rhythms of maternal blood pressure remains unclear. Moreover, the effects of maternal immune system dysregulation during pregnancy on placental clock genes have not been established.

In this study, we used the gravid rat as an experimental model to determine the impact of exposure to unmethylated CpG DNA during pregnancy on circadian patterns of maternal cardiovascular parameters, expression of clock genes in the placenta, and the interactive effects of these consequences on perinatal outcomes. We treated rats with ODN2395 (type of CpG ODN) as an immune trigger and synthetic stimulant of TLR9. Previous studies have shown that exposure to CpG ODN during pregnancy induced adverse maternal cardiovascular and perinatal outcomes in rats (37–39). We hypothesized that maternal treatment with CpG ODN during rat pregnancy would elicit aberrant inflammatory responses, disrupt maternal blood pressure circadian rhythms, and dysregulate the placental molecular clock machinery, mediating adverse fetoplacental outcomes.

## METHODS

All protocols were approved by the Institutional Animal Care and Use Committees (IACUC) of the University of North Texas Health Science Center (IACUC-2020-032) and Loma Linda University (IACUC-22-003) performed in accordance with the Guide for the Care and Use of Laboratory Animals of the National Institutes of Health. All investigators followed the ethical principles outlined in the ARRIVE guidelines.

### Animals

Male (body weight and age on arrival: 400 g and 12-15 weeks) and virgin female (body weight and age on arrival: 200 g and 9-11 weeks) Sprague-Dawley rats (Envigo, Indianapolis, IN and Houston, TX, USA) were single-housed under 12 h:12 h light/dark cycles (lights on, 0700 h; lights off, 1900 h) in a temperature and humidity-controlled environment. Animals were provided standard laboratory chow and water ad libitum. Male rats were only used for breeding purposes. Following one week of acclimatization to the animal facilities, female rats were familiarized with handling and vaginal smears before any experimentation, and a normal estrous cycle was determined using vaginal cytology. For pregnancy studies, female rats were mated in-house (pair mating) overnight, and vaginal smears were assessed the following morning for presence of spermatozoa. Gestational day (GD) 1 (term = 22-23 days) was designated as the morning on which spermatozoa were observed in vaginal smears, and daily body weights were recorded throughout gestation. All experiments were performed when rats were 12-24 weeks old. In total, 47 female and 12 male rats were used for these studies.

### Experimental design and timeline

Two studies were conducted and included two separate cohorts of animals (Figure 1). The purpose of the first study (Cohort 1) was to determine the impact of repeated exposure to CpG ODN, on maternal hemodynamics, circadian rhythms of maternal blood pressure and heart rate, plasma cytokine concentrations, and fetoplacental biometrics. The purpose of the second study (Cohort 2) was to determine acute maternal systemic inflammatory responses to a single dose of CpG ODN and the impact of pregnancy on these responses. Placental inflammation and clock gene expression was also measured in placentas from rats exposed to a single dose of CpG ODN to determine placental responses in the absence of chronic systemic responses to this treatment. These studies were conducted only in female rats because the focus of this research is on maternal physiology during pregnancy, which is a female-specific condition.

**Figure 1.**
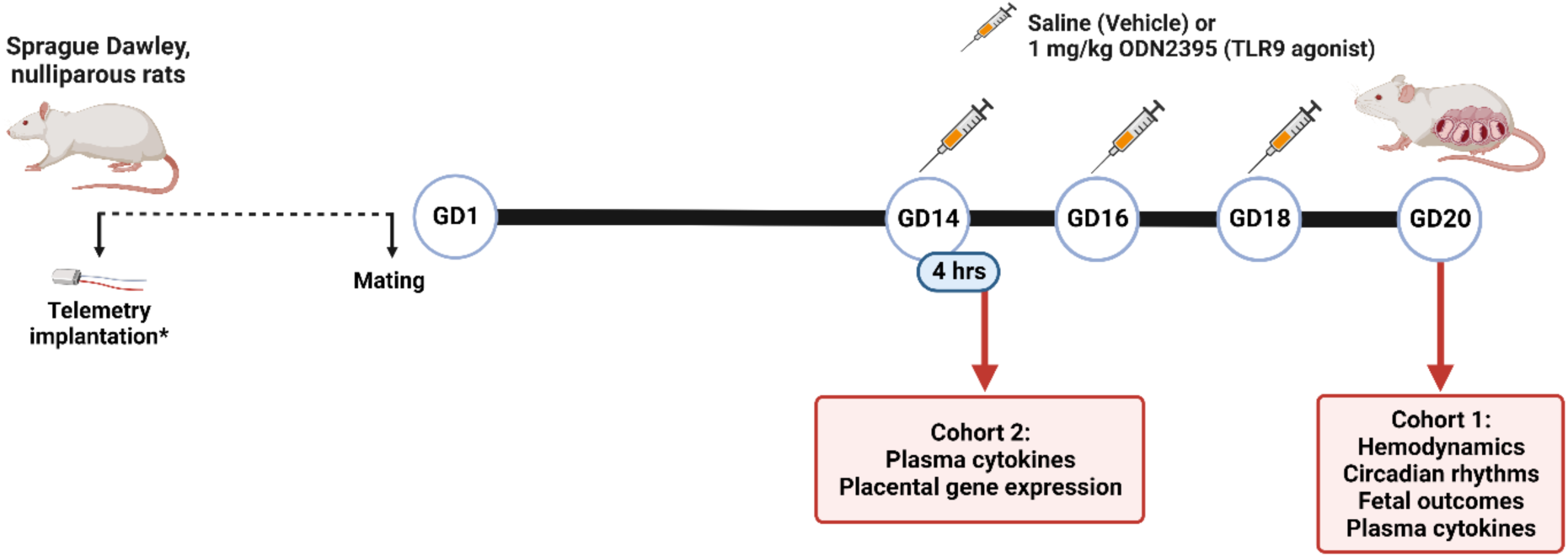
Experimental design and timeline. Two cohorts were included in the experimental design. Rats assigned to Cohort 1 were virgin, non-pregnant females implanted with telemeters prior to mating, followed by repeated intraperitoneal (IP) injections of Saline (vehicle) or 1 mg/kg ODN2395 during the third trimester of their pregnancy on gestational day (GD) 14, GD16, and GD18, with euthanasia on GD20 to assess maternal hemodynamics, circadian rhythms of maternal blood pressure and heart rate, fetal outcomes, and maternal plasma cytokines. Cohort 2 included pregnant females at GD14 and age-matched virgin, non-pregnant females. Rats assigned to Cohort 2 were intraperitoneally (IP) administered Saline (vehicle) or 1 mg/kg ODN2395 and euthanized 4 hours after injections to assess maternal plasma cytokines and placental gene expression. TLR: Toll-like receptor 9; *, telemeters were only implanted in Cohort 1.

### Animal treatments

Female rats were given intraperitoneal injections of 300 µl of 0.9% saline (vehicle) or CpG ODN (ODN2395,1 mg/kg body weight; InvivoGen, Cat. # tlrl-2395; TLR9 agonist). ODN2395 has unmethylated CpG sequences (5’-TCGTCGTTTT<u>CGGCGC:GCGCCG</u>-3’, palindrome is underlined) and is designed with a phosphorothioate-modified backbone for increased resistance to nucleases resulting in a greater half-life of approximately 1 hour (40). The dose of ODN2395 used in treatments was based on the manufacturer’s recommended range and preliminary studies. Notably, rodent TLR9 has rhythmic expression with greatest expression levels and activity during the wake phase (41). Thus, injections were given in the afternoon between 1600-1800 h (just prior to wake phase). Cohort 1 included pregnant rats that were treated on GD14, GD16, and GD18 and euthanized on GD20 as previously described (37, 38). Cohort 2 included pregnant rats that were treated on GD14 and age-matched non-pregnant rats. Rats assigned to Cohort 2 were euthanized four hours after treatment.

### Radiotelemetry implantation and maternal hemodynamic measurements

To assess blood pressure and heart rate in freely moving conscious rats, non-pregnant female rats (Cohort 1) were implanted with an abdominal aortic catheter attached to a HD-S10 radiotelemeter transmitter (Data Sciences International, St. Paul, MN). Radiotelemeters were turned on one week following implantation, and maternal systolic blood pressure (SBP), diastolic blood pressure (DBP), mean arterial pressure (MAP), and heart rate (HR) were measured during a 10-s sampling period (500 Hz) and were averaged and recorded every 5 min using Ponemah software (Data Sciences International, St. Paul, MN). Three days of baseline hemodynamics were recorded prior to mating, and telemetry measurements continued throughout gestation. Data are first summarized as either a 12-hour average of dark cycle (wake phase) or 12-hour average of light cycle (sleep phase) and are presented as absolute values (i.e., mmHg or beats per minute). Blood pressure and heart rate tracings were used for circadian rhythm analysis as described below.

### Euthanasia and tissue harvest

Rats were anesthetized with isoflurane (5% for induction, 3% for maintenance) and euthanized by isoflurane overdose followed by bilateral thoracotomy and removal of their hearts. Whole blood was collected from the inferior vena cava when rats were under a deep plane of anesthesia. Following euthanasia, plasma was isolated from whole blood using EDTA-coated collection tubes (BD, Cat. # 367856) centrifuged at 1700 x g for 15 min at 4°C. Freshly isolated plasma was snap frozen in liquid nitrogen and stored at −80°C until further analysis. The left and right uterine horns containing corresponding fetuses and placentas were excised, and fetoplacental biometrics were recorded prior to fetal euthanasia via decapitation. Maternal deciduae were removed from placentas prior to snap freezing in liquid nitrogen and storing at −80°C for subsequent analysis.

### Plasma cytokine and chemokine analyses

Plasma cytokines and chemokines were assessed using a magnetic bead panel (Bio-Rad, Cat. #171K1001M). Plasma samples were diluted 1:4 in assay buffer prior to running the assay according to manufacturer’s instructions. All samples and standards were run in duplicate, and plates were read on the Bio-Plex^®^ MAGPIX™ Multiplex Reader (Bio-Rad, Hercules, CA) using xPONENT® software version 4.3 (Luminex Corporation, Austin, TX). The following analytes were measured: proinflammatory cytokines (TNF-α, IL-6, IL-1β, IL-1α, IL-18, IL-12p70, IL-17A, IFNγ, IL-7), anti-inflammatory cytokines (IL-4, IL-5, IL-13, IL-10, IL-2), and chemoattractants (GRO/KC, MCP-1, MIP-1α, MIP-3α, and RANTES).

### Extraction of RNA from placentas

RNA was isolated from GD14 placentas after removal of maternal decidua. To extract RNA, placentas were digested in 1 ml of TRIzol Reagent (Invitrogen, Cat # 15596018) per 100 mg tissue and homogenized on ice using a Qsonica sonicator (3 sets of 3s pulses with 10 min rest on ice between pulsing). Homogenized placentas were treated with 200 µl chloroform (Sigma, Cat # C2432) per 1 ml TRIzol used during digestion, vortexed, and allowed to rest at room temperature for 2 minutes prior to centrifuging at 12,000 x g for 15 minutes at 4°C. RNA was precipitated from the aqueous phase using 500 µl isopropyl alcohol (Sigma, Cat # I9516) per 1 ml TRIzol used during homogenization. After allowing RNA to precipitate for 10 minutes at room temperature, samples were centrifuged at 12,000 x g for 10 minutes at 4°C. Following centrifugation, the supernatant was removed, and RNA pellet was washed with 75% ethanol, dried, and suspended in RNase-free water. RNA was dissolved by incubating samples at 60°C for 10 min. The purity and quantity of total RNA isolated from each sample was determined using a Nanodrop spectrophotometer (Thermo Scientific). All samples had a 260/280 ratio of 1.98-2.07 and were diluted to 50 ng/µl RNA using RNase-free water.

### Synthesis of cDNA

RNA was reverse transcribed to cDNA using Sensiscript RT Kit reagents (Qiagen, cat # 205213) with RiboGuard RNase inhibitor (Lucigen, Cat # E0126-40D7) and oligo-dT primers (Qiagen, cat # 79237). Each 20 µl reaction included: 2 µl 10X RT Buffer, 2 µl dNTP mix, 2 µl oligo-dT primers, 0.25 µl RNase inhibitor (40 U/µl), 1 µl Sensiscript RT, 8.75 µl RNase-free water, 4 µl of 50 ng/µl RNA. cDNA was synthesized using a T100 thermocycler (Bio-RAD) set at 37°C for 1 hr. Synthesized cDNA was stored at −20°C until subsequent analysis.

### Quantitative real-time polymerase chain reaction (qRT-PCR)

The expression of rat placenta proinflammatory cytokine (*Tnfα, Il6, Il1β*), clock (*Clock, Bmal1, Cry1, Per1, Per2, Per3*), and reference (*Ppia* and *Sdha*) genes was determined using qRT-PCR. Primer sequences were identified from previous studies (Supplementary Table S1)) and purchased from Integrated DNA Technologies, Inc (IDT). Reactions consisted of 7.5 µl iQ SYBR Green Supermix (Bio-Rad, Cat # 1708882), 1.2 µl primer mix (50 µl of 100 mM forward primer, 50 µl of 100 mM reverse primer, 100 µl RNase-free water), 1.8 µl cDNA, and 4.5 µl RNase-free water. Non-template control (NTC) reactions were included for each primer set. Each sample and NTC were ran in duplicate for target and reference genes. qRT-PCR was performed on a CFX96 Real-Time PCR Detection System with CFX Maestro Software v 2.3 (Bio-Rad). Cycling parameters were initial denaturation at 95°C for 3 min followed by 40 cycles of 95°C for 10 s and 60°C for 1 min. At the end of the cycling, melt curves were generated by 5 s interval increases in 5°C from 65°C to 95°C. Primer specificity was determined by the presence of a single melt peak in samples and no generated melt peaks in NTC. Gene expression was analyzed using the 2^-ΔΔCT^ method. *Ppia* was chosen for normalization based on its stable expression across all samples.

### Sample size determination, data, and statistical analyses

The smallest sample size needed to determine differences in blood pressure responses between rats treated with ODN2395 or vehicle was estimated using power analysis based on our previous and preliminary observations. The sample size was selected with a goal to achieve a power of 0.80 to 0.85 with a probability of a Type I error of 0.05.

Unpaired t tests were used to determine group differences in plasma cytokine concentrations and fetoplacental outcomes. Hemodynamic responses over the treatment window (GD14-GD19) in saline and ODN2395-treated rats were analyzed by fitting a mixed model with Geisser-Greenhouse correction as implemented in GraphPrism (version 9.2, GraphPad Software, Inc.). This mixed model uses a compound symmetry covariance matrix and is fit using Restricted Maximum Likelihood (REML). We used this analysis instead of ANOVA with repeated measures because ANOVA cannot handle missing values. In our data set, values were missing completely at random due to occasional connectivity issues with the radiotelemeters. To evaluate the effect of advancing gestational age on hemodynamic responses within the treatment group (GD15, GD16, GD17, GD18, GD19 vs. GD14), we performed multiple comparisons (Dunnett’s test) and calculated multiplicity adjusted *p*-values for each comparison. Outliers were determined using ROUT (**Ro**bust regression and **Out**lier removal, ROUT coefficient Q = 1%), and normal distributions were tested using Shapiro-Wilk test (GraphPrism). Data determined to have non-Gaussian distributions were normalized using log transformation before applying parametric statistics. For clarity, raw data are presented in figures. The relationship between placental clock gene expression and inflammatory gene expression was determined using Spearman correlation analysis. The significance level was set to α = 0.05 and *p* ≤ 0.05 was considered significant. Values are expressed as mean ± standard deviation (SD) unless otherwise noted.

Multiple linear regression analysis of fetoplacental outcomes was incorporated to examine the effect of covariates (number of resorptions, average placental weight, and litter size) on the main outcome of fetal weight (R statistical software version 4.0.2). Multivariate outliers were detected and removed by Median Absolute Distance (MAD) using principal component analysis (PCA) on fetal biometrics. Sample measures with PC scores ≥ 2 SDs from median PC score were deemed outliers. After outlier removal, multicollinearity was detected between the independent variables: number of resorptions and litter size. To remove multicollinearity, we assessed 3 multiple linear models and determined best fit to our data using the Akaike’s Information Criterion (AIC): (1) average fetal weight as a function of treatment, litter size, and average placental weight, (2) average fetal weight as a function of treatment, average placental weight, and number of resorptions, and (3) average fetal weight as a function of treatment, average placental weight, and ratio of resorptions within a litter. Model 2, average fetal weight as a function of treatment, average placental weight, and number of resorptions, displayed the best fit (AIC = −50.3, Supplementary Table S2) and was therefore used to draw conclusions.

We used Lomb-Scargle periodogram to detect and analyze circadian rhythmicity and characteristics of the identified rhythms in blood pressure and heart rate time series data. Lomb-Scargle periodogram analysis was performed on continuous 24-h raw data with a resolution of 5-min bins for each gestational day using R (DiscoRhythm package (42), R statistical software version 4.0.2, R Core Team 2020) after outlier detection and removal by anomaly detection (Anomalize package for R (43)). DiscoRhythm determines a common period across multiple parallel time series (i.e., able to find a common period given multiple subjects monitored at the same time (42). Rhythmicity was determined for each treatment window, which was defined as point of injection_n_ to point of injection_n+1_, per rat per treatment on 24 h cycle. A *p*-value ≤ 0.05 indicates significant period (i.e., circadian rhythm) detected. The following rhythmic parameters were computed: acrophase and amplitude. Acrophase denotes the time of maximum value (peak) and is expressed in Zeitgeber time (0-12 = sleep phase, 0700 h – 1900 h; 12-24 = wake phase, 1900 h-0700 h), while amplitude denotes the magnitude of the peak at given acrophase and is expressed in native units (mmHg for blood pressure and BPM for heart rate). We selected the Lomb-Scargle periodogram for rhythmicity detection because it allows the analysis of data sets with missing values (e.g., due to transient signal loss) and is considered to have a better detection efficiency and accuracy in the presence of noise, while it avoids bias that could arise from replacement of missing data by interpolation techniques (44). Specifically, Lomb-Scargle periodogram has been suggested as a suitable method to study hemodynamic variable rhythms in telemetrical time-series recorded from living animals (45).

## RESULTS

### Maternal blood pressure and heart rate responses during sleep and wake cycles and their circadian rhythmicity in rats exposed to CpG ODN during pregnancy

There were no differences in pre-pregnancy sleep and wake SBP, DBP, MAP, or HR between rats assigned to saline vs. ODN2395 groups (*p* > 0.05, Supplementary Table S3). To determine the effects of repeated exposure to treatment with ODN2395 on maternal hemodynamics, we used mixed effect analysis on data collected using radiotelemetry during the treatment window (GD14 – GD19, GD14: before treatment started, GD19: after the last treatment) in saline-treated and ODN2395-treated groups. We found that the effect of treatment (responses in saline-treated rats minus responses to ODN2395) was consistent across the treatment window (GD14 – GD19) for SBP (SBP, Figure 2A-B), DBP (Figure 2C-D), MAP (Figure 2E-F) or HR (Figure 2G-H) in the sleep phase (time x treatment interaction, *p ≥* 0.33) or wake phase (time x treatment interaction, *p ≥* 0.21). Multiple comparisons tests with adjustments for multiplicity were used to evaluate the effect of advancing gestational age on hemodynamic responses within each treatment group. Sleep DBP was lower on GD18 (*p* = 0.04, mean difference ± SE: 4.5 ± 1.2 mmHg) and GD19 (*p* = 0.003, mean difference ± SE: 7.0 ± 1.0 mmHg) compared to GD14 in saline-treated rats. In a similar manner, sleep MAP was lower on GD19 (*p* = 0.02, mean difference ± SE: 6.5 ± 1.5 mmHg) compared to GD14 in saline-treated rats. However, reduction in sleep maternal blood pressure at the end of pregnancy was not seen in ODN2395-treated rats (*p* > 0.05). These data suggest that blood pressure recorded during the sleep phase decreases with advancing gestational age in healthy pregnant rats but not in rats treated with ODN2395.

**Figure 2.**
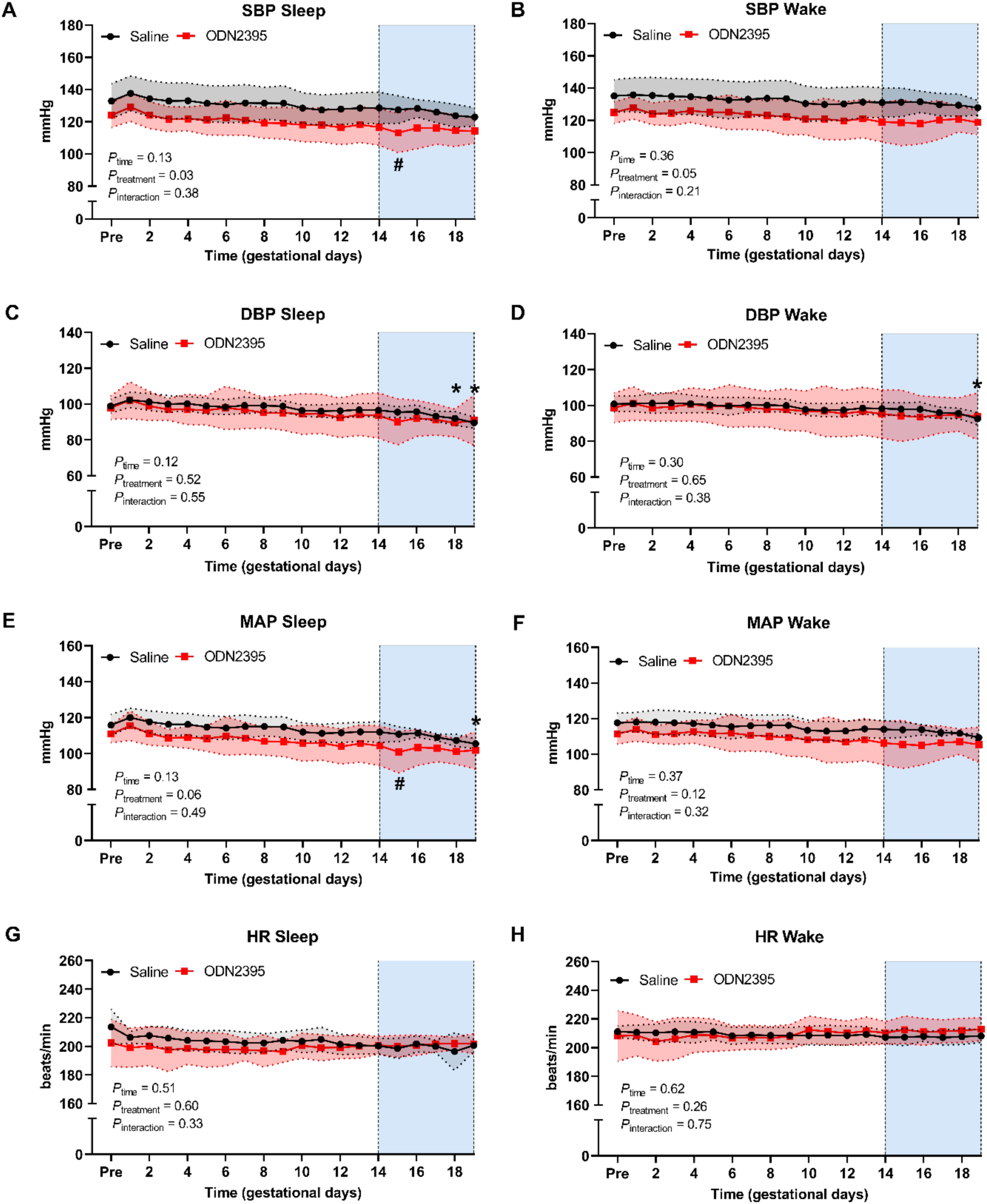
Temporal changes in maternal blood pressure and heart rate responses during sleep and wake cycles before and throughout gestation in rats treated with ODN2395 or saline vehicle. A, B) Systolic blood pressure, C, D) diastolic blood pressure, E, F) mean arterial pressure, and G, H) heart rate during sleep (A, C, E, G) and wake (B, D, F, H) cycles. Intraperitoneal injections of ODN2395 or saline (vehicle) were given on gestational day (GD)14, 16, and 18. Shaded vertical region represents treatment window (GD14-GD19). Data within the treatment window were analyzed by fitting a mixed model with Geisser-Greenhouse correction. Multiple comparisons to assess simple effects were performed using Dunnett’s test (responses on GD15, GD16, GD17, GD18, GD19 vs. GD14 within each treatment group) and multiplicity adjusted *p*-values for each comparison were calculated. Simple effects: * = *p* < 0.05 vs. GD14 in saline group, # = *p* < 0.05 vs. GD14 in ODN2395-treated group. Values are presented as mean ± SD; n = 7/group. SBP = systolic blood pressure, DBP = diastolic blood pressure, MAP = mean arterial pressure, HR = heart rate, PRE = three-day average of values prior to mating.

Sleep SBP measured after the first treatment of ODN2395 (GD15) was lower compared to SBP before treatment, on GD14 (*p* = 0.009, mean difference ± SE: 3.5 ± 0.69 mmHg). This reduction in SBP contributed to a decrease in sleep MAP on GD15 compared to GD14 in the ODN2395 group (*p* = 0.008, mean difference ± SE: 3.5 ± 0.70 mmHg). After GD15, sleep blood pressure of ODN2395-treated rats returned to pre-treatment values and remained unchanged throughout the rest of the treatment window (GD16, GD17, GD18, GD19 vs. GD14, *p* > 0.05). These data suggest that SBP and MAP recorded during the sleep phase were reduced in response to the first dose of ODN2395 and returned to pre-treatment values despite subsequent repeated exposure to ODN2395.

A significant circadian rhythm was detected for all hemodynamic variables over the treatment window (Figure 3). Circadian rhythm was lost for both SBP and DBP after the first ODN2395 treatment on GD14 but recovered and was unaffected after the second treatment with ODN2395 on GD16 (Figure 3A, Table 1). After the last treatment on GD18, DBP lost circadian rhythmicity (Figure 3A, Table 1). The amplitude of HR was magnified in the ODN2395 group, and the period of HR was slightly later, but otherwise HR was unaffected (Figure 3B, Table 1).

**Figure 3.**
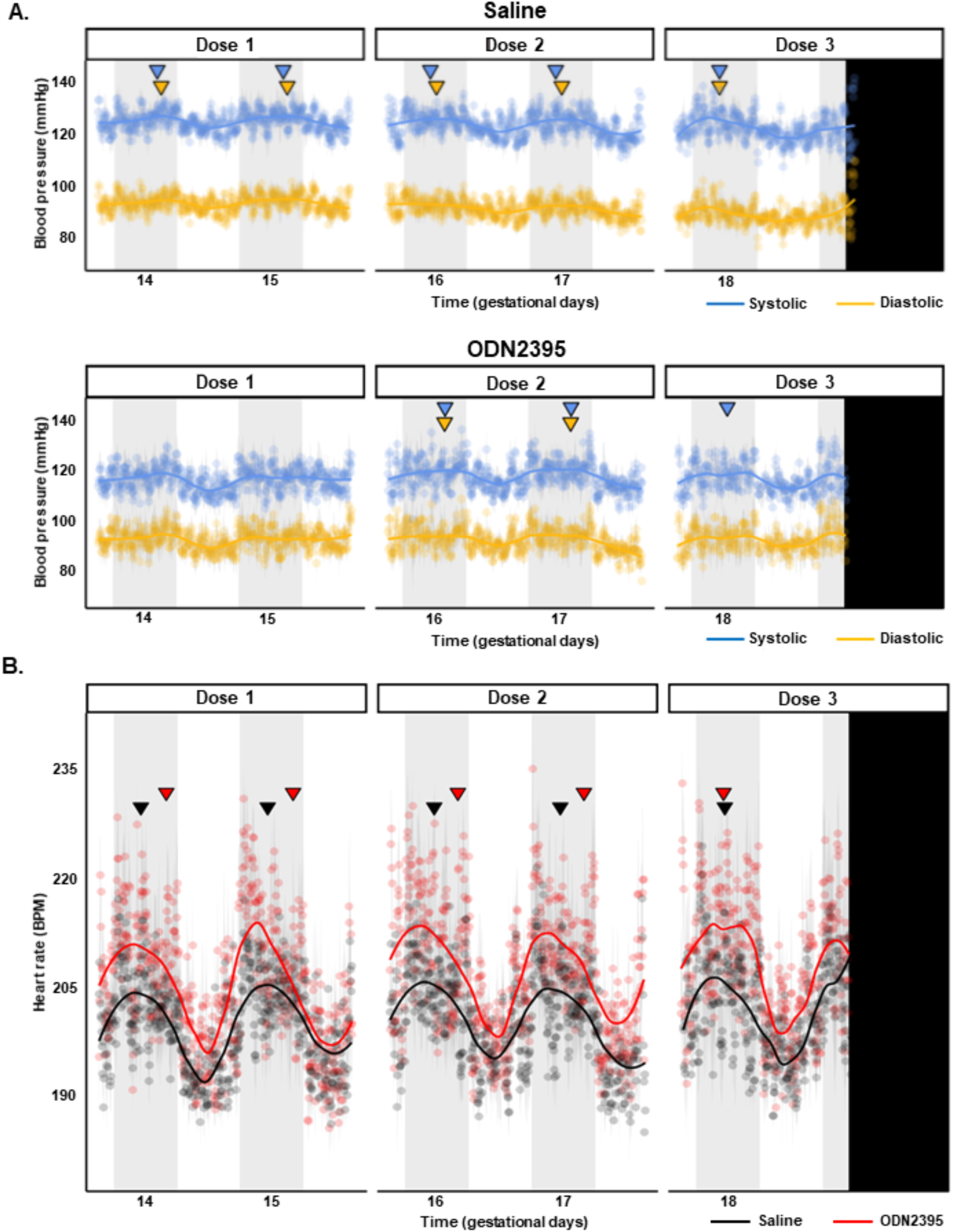
Circadian rhythms of maternal blood pressure and heart rate during late gestation in rats treated with ODN2395 or saline vehicle. A) Mean systolic (blue) and diastolic (gold) blood pressures during treatment window (Gestational day, GD, 14 – GD19) for saline (top panel) and ODN2395 (bottom panel) cohorts. B) Mean heart rate during treatment window (GD14 – GD19) for saline (black) and ODN2395 (red) cohorts. Blue (systolic) and gold (diastolic) inverted triangles (panel A) represent significant acrophases of respective blood pressures, while black (saline) and red (ODN2395) inverted triangles represent significant acrophases of heart rate circadian rhythm. Acrophases denote the time of maximum value (peak) and were calculated by Lomb-Scargle periodogram analysis performed on continuous 24-h raw data with a resolution of 5-min bins. Lack of inverted triangle indicates no significant circadian rhythm detected. Vertical black bar denotes end of measurement window (midnight GD20). Grey vertical bars represent dark hours (active phase). Shaded area flanking data points represents standard error; n = 7/group. Trendlines estimated by loess smoothing specifying an alpha of 0.4. BPM = beats per minute.

**Table 1.** Lomb-Scargle periodogram analysis of maternal blood pressure and heart rate circadian rhythms.

|  | Saline (n = 9) |  |  | ODN2395 (n = 7) |  |  |
| --- | --- | --- | --- | --- | --- | --- |
|  | SBP | DBP | HR | SBP | DBP | HR |
| **Acrophase (ZT)** |  |  |  |  |  |  |
| GD 14-GD 16 | 20:15* | 21:00* | 16:55* | **20:12** | **20:28** | 21:44* |
| GD 16-GD 18 | 17:00* | 18:11* | 17:13* | 20:05* | 19:58* | 21:44* |
| GD 18-GD 20 | 17:00* | 17:00* | 17:14* | 18:44* | **17:00** | 17:00* |
| **Amplitude (mmHg/BPM)** |  |  |  |  |  |  |
| GD 14-GD 16 | 130.12* | 97.23* | 206.06* | **117.93** | **93.68** | 213.31* |
| GD 16-GD 18 | 129.48* | 95.36* | 205.83* | 119.61* | 93.79* | 211.93* |
| GD 18-GD 20 | 130.44* | 94.42* | 207.84* | 118.43* | **93.22** | 216.69* |

Acrophase in Zeitgeber time (0-12 = sleep phase, 0700 h – 1900 h; 12-24 = wake phase, 1900 h-0700 h) and denotes the time of maximum value (peak). Amplitude expressed in native units (mmHg for BP and BPM for HR) and denotes the magnitude of the peak at given acrophase. \* = significant periodicity detected ( $p < 0.05$ ). Bolded text = no significant periodicity detected, representing a loss of circadian rhythmicity. SBP = systolic blood pressure, DBP = diastolic blood pressure, HR = heart rate, BPM = beats per minute.

### Fetoplacental biometrics in response to gestational exposure to CpG ODN

Univariate analysis showed no differences in mean fetal weights (Saline: 2.25 ± 0.18 g, ODN2395: 2.28 ± 0.09 g, *p* = 0.67, Figure 4A), placental weights (Saline: 0.49 ± 0.04 g, ODN2395: 0.44 ± 0.06 g, *p* = 0.06, Figure 4B), fetoplacental ratio (Saline: 4.69 ± 0.45, ODN2395: 5.20 ± 0.59, *p* = 0.07, Figure 4C), litter size (Saline: 14.78 ± 1.56, ODN2395: 14.86 ± 2.41; *p* = 0.94, unpaired t-test) or number of resorptions per litter (Saline: 2.22 ± 2.17, ODN2395: 0.86 ± 90; *p* = 0.14, unpaired t-test) on GD20. Fetal weights vary with litter size and thus, previous toxicology studies suggested that litter size and number of resorptions should be considered when the effect of a test substance on fetal weights is measured (46). Therefore, in addition to univariate analysis for mean comparisons between groups, we assessed the effects of ODN2395 treatment on the relationship between perinatal outcomes, namely fetal weights, placental weights, and number of resorptions. Notably, we found increases in the number of resorptions was disproportionately associated with reduced fetal weights and placental weights in ODN2395-administered dams compared to saline-administered dams (Figure 4D). Treatment with ODN2395 amplified the relationship of reduced fetal weights with increased resorptions in the litter, promoted overall lower placental weight when resorptions are present, and reduced the association of larger placentas and increased fetal weight (Figure 4D; weighted multiple linear regression, treatment x resorption & treatment x placental weight x resorptions, *p* = 0.02, Supplementary Table S4). Collectively, these data suggest that repeated exposure to CpG ODN resulted in interactive effects on pregnancy dynamics that contributed to reduced fetal weight (Figure 4D, Supplementary Table S4). Of note, treatment with CpG ODN did not impact maternal weight gain during pregnancy (*p* = 0.54, Supplementary Figure S1).

**Figure 4.**
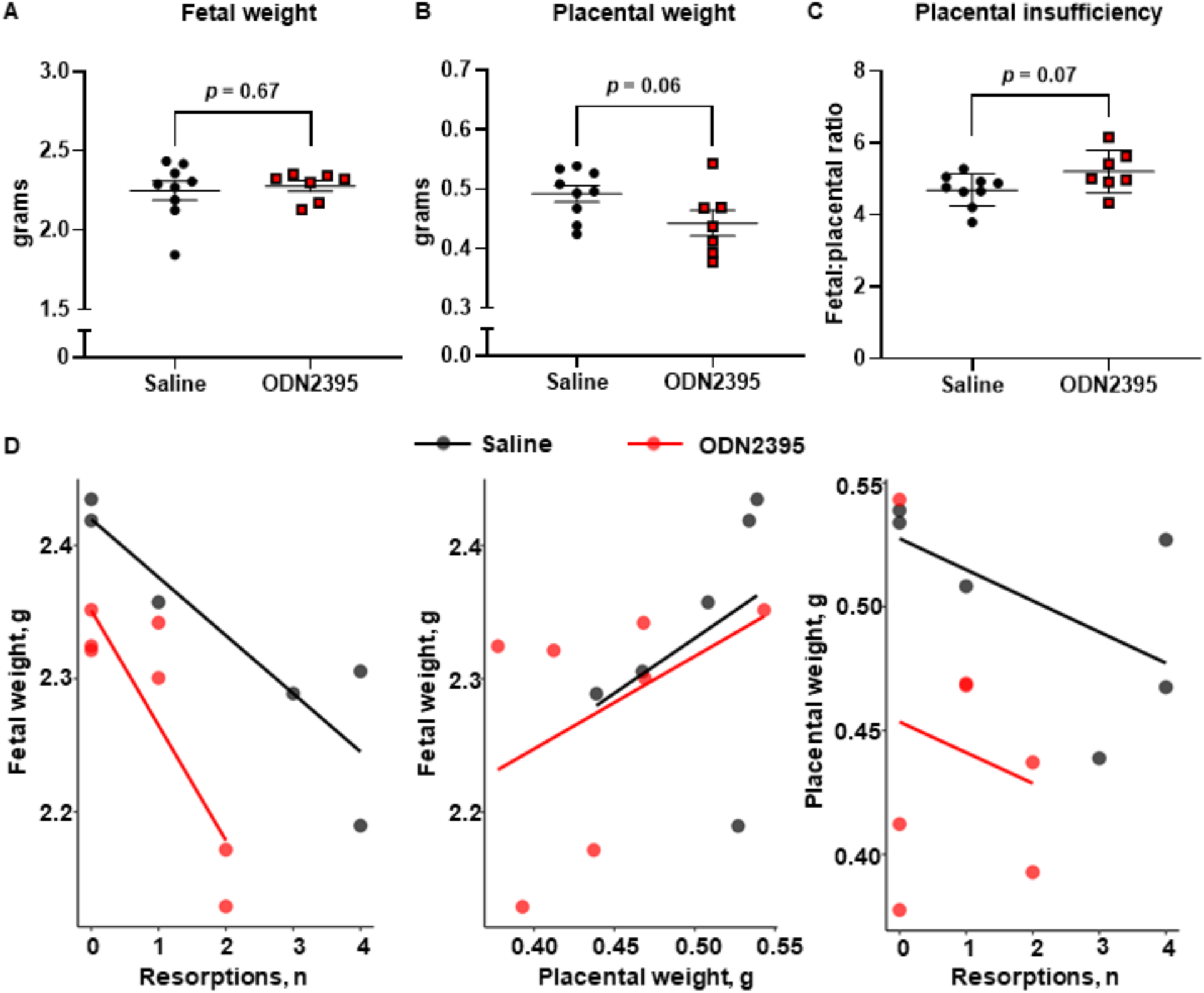
Effects of ODN2395 on fetal and placental biometrics and fetoplacental dynamics on gestational day 20. Individual and mean A) fetal weights (g) per litter, B) placental weights (g) per litter, C) fetal to placental weight ratio per litter, and D) scatterplots representing correlations of average fetal weight (g) with number of resorptions (left panel), average fetal weight (g) with average placental weight (g, middle panel), and average placental weight (g) with number of resorptions (right panel). Each dot represents the average from a single litter (n = 7-9/group). A-C) Values are presented as mean ± SD, unpaired t-test; D) Trendlines (Saline = black, ODN2395 = red) represent the linear relationships within the data determined by the multiple linear regression.

### Inflammatory cytokine and chemoattractant concentrations in maternal circulation after exposure to CpG ODN

Pregnant rats treated with ODN2395 in late pregnancy (GD14, 16, 18) had reduced plasma concentrations of proinflammatory cytokines and monocyte chemoattractants at GD20 (Figure 5, Supplementary Table S5). Remarkably, maternal plasma concentrations of IL-1α and IL-17A were reduced in ODN2395-treated dams compared to controls at GD20 (*p* = 0.03, Figure 5A, Supplementary Table S5). Additionally, maternal plasma concentrations of chemoattractants MCP-1 and MIP-3α were reduced in the ODN2395 group compared to control at GD20 (*p* ≤ 0.05, Figure 5B, Supplementary Table S5).

**Figure 5.**
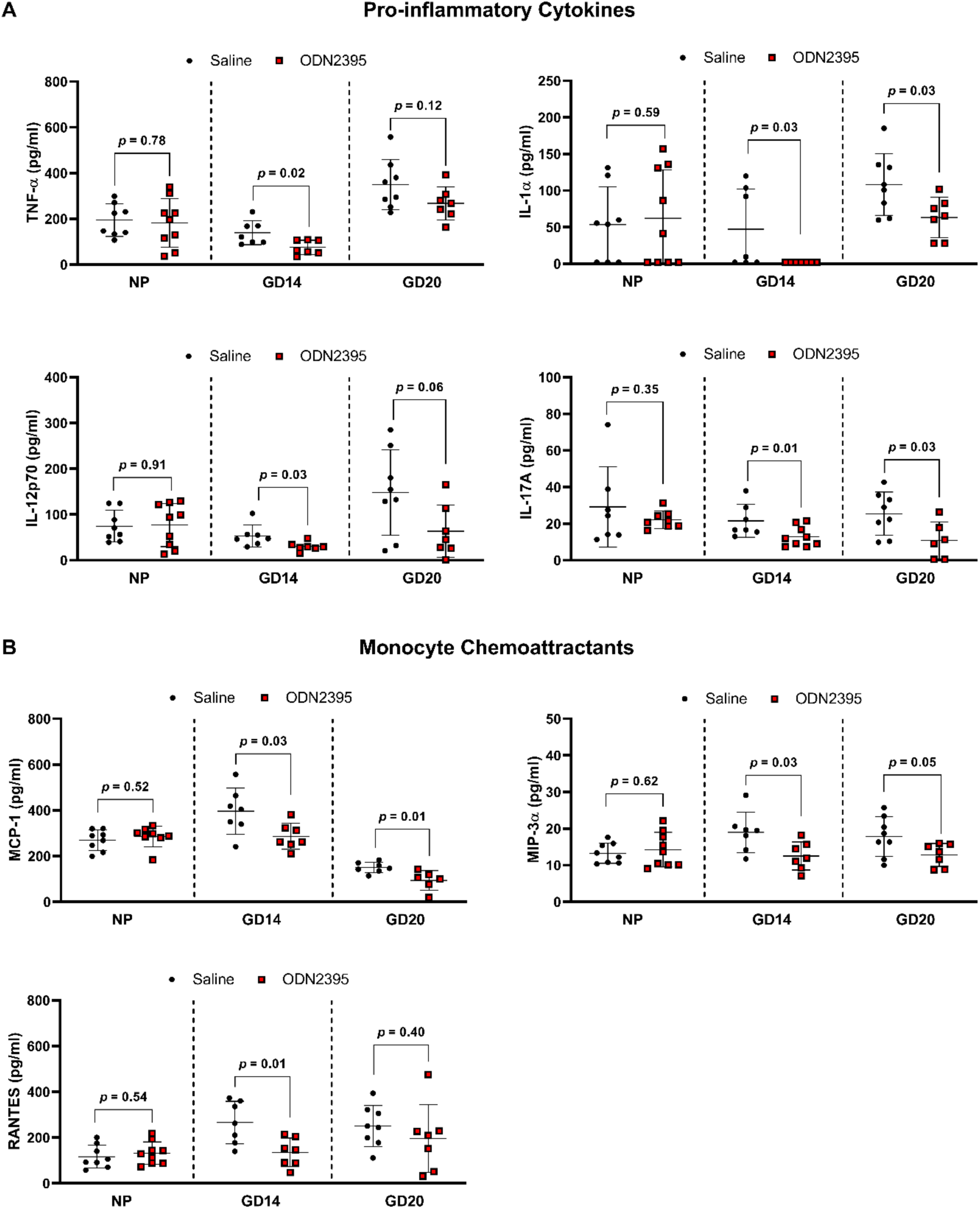
Effects of a single and repeated exposures to ODN2395 on circulating proinflammatory cytokines and monocyte chemoattractants in non-pregnant and pregnant rats. A) Proinflammatory cytokines and B) monocyte chemoattractants in plasma of non-pregnant, GD14 dams (Cohort 1), and GD20 dams (Cohort 2) intraperitoneally administered ODN2395 or saline vehicle. Non-pregnant and GD14 dams received a single IP injection while GD20 dams received three sequential IP injections of ODN2395 or saline vehicle. Values are presented as mean ± SD (n = 7-9/group) and were analyzed by unpaired t-test. Black circles = Saline; Red boxes = ODN2395. NP = non-pregnant, GD = gestational day, TNF-α = Tumor necrosis factor-alpha, IL-1 α = Interleukin-1 alpha, IL-12p70 = bioactive dimer of Interleukin-12 p35 and p40 subunits, IL-17A = Interleukin-17A, MCP-1= Monocyte chemoattractant protein-1, MIP-3α = Macrophage inflammatory protein-3 alpha, RANTES = Regulated on Activation, Normal T Cell Expressed and Secreted.

Since we observed a suppressed inflammatory response to repeated treatments of ODN2395 during late pregnancy, we assessed the impact of a single dose of ODN2395 on the maternal proinflammatory profile at GD14. We quantified plasma levels of maternal cytokines and monocyte chemoattractants four hours after a single dose of ODN2395. Similar to our results in GD20 dams, our data demonstrated ODN2395 exposure during pregnancy reduced plasma proinflammatory cytokine and monocyte chemoattractant concentrations at GD14 (Figure 5, Supplementary Table S5). Specifically, a single dose of ODN2395 during pregnancy suppressed plasma concentrations of proinflammatory cytokines TNF-α, IL-1β, IL-1α, IL-12p70, IL-17A, and IL-7 (*p* ≤ 0.03, Figure 5A, Supplementary Table S5) and monocyte chemoattractants MCP-1, MIP-3α, and RANTES (*p* ≤ 0.03, Figure 5B, Supplementary Table S5). There was no effect of repeated ODN2395 exposure on circulating anti-inflammatory cytokines at GD20 (Supplementary Table S5). However, a single dose of ODN2395 at GD14 reduced anti-inflammatory cytokines IL-5 and IL-2 (*p* ≤ 0.04, Supplementary Table S5). Similar reductions in circulating maternal anti-inflammatory cytokines IL-4 and IL-10 were also observed in the ODN2395-treated rats compared to controls, but these differences did not reach statistical significance (*p* = 0.06 and 0.08, respectively; Supplementary Table S5). Of note, ODN2395-mediated reductions in circulating cytokines and monocyte chemoattractants were pregnancy-specific, as evidenced by no effect of ODN2395 exposure on the systemic immune profile in virgin, non-pregnant females (*p* ≥ 0.34, Figure 5 and Supplementary Table S5).

### Placental inflammation and clock gene expression after exposure to CpG ODN

Placentas from dams treated with a single dose of ODN2395 on GD14 had greater expression of proinflammatory *Tnfα* compared to placentas from saline-treated dams, but there were no group differences in expression of proinflammatory *Il6* or *Il1β* (Figure 6). In addition, placental expression of CLOCK and BMAL-1 target genes *Per2* and *Per3* were increased after a single treatment with ODN2395 (*p* ≤ 0.05, Figure 7). No effects of ODN2395 exposure were observed in placental gene expression of *Clock* or *Bmal1* (*p* ≥ 0.13, Figure 7) or on CLOCK and BMAL-1 target genes *Cry1* or *Per1 (p* ≥ 0.15, Figure 7). There was a negative correlation between placental *Tnfα* and *Bmal1* (r = −0.63, *p* = 0.02, Figure 8) and a positive correlation of placental *Tnfα* and *Per3* (r = 0.56, *p* = 0.06, Figure 8). There were no significant correlations between placental *Tnfα* expression and the expression of *Clock*, *Bmal1*, *Cry1*, or *Per2* (*p* ≥ 0.27, Figure 8).

**Figure 6.**
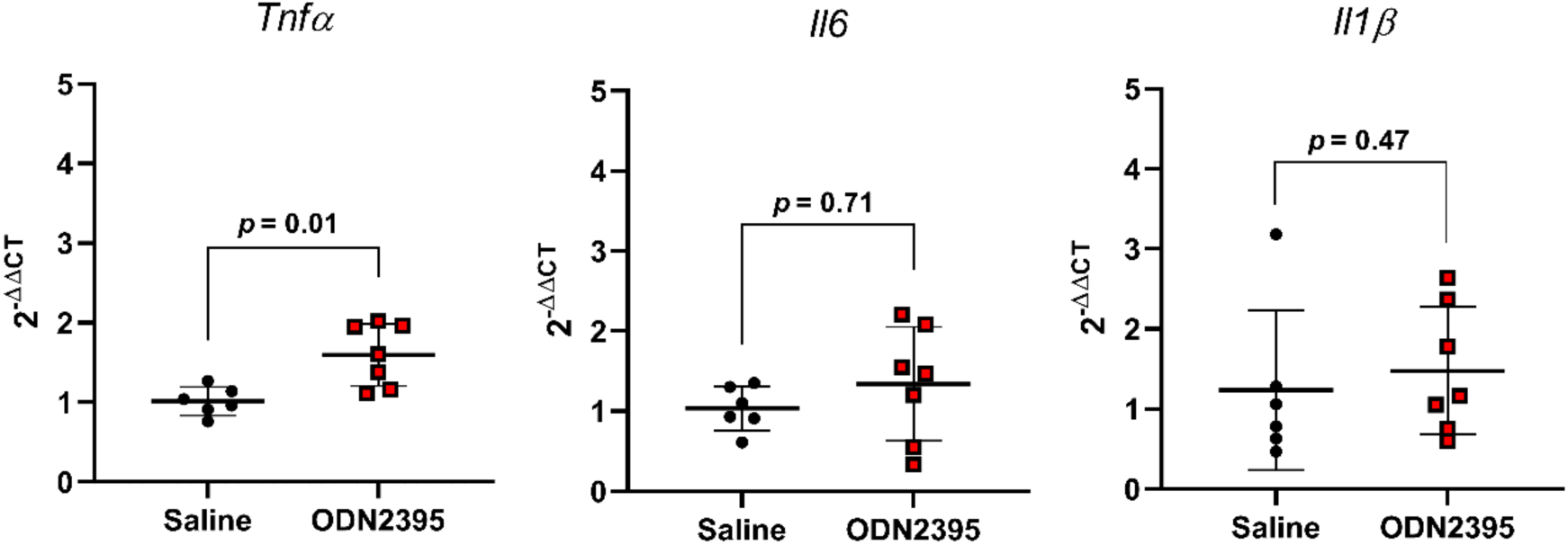
Effect of single exposure to ODN2395 on proinflammatory cytokine gene expression in rat placentas on gestational day 14. Placental gene expression of tumor necrosis factor alpha (*Tnfα*), Interleukin-6 (*Il6*), and Interleukin-1β (*Il1β*) four hours after administration of ODN2395 or saline vehicle. Values are presented as mean ± SD (n = 6-7/group) and were analyzed by unpaired t-test.

**Figure 7.**
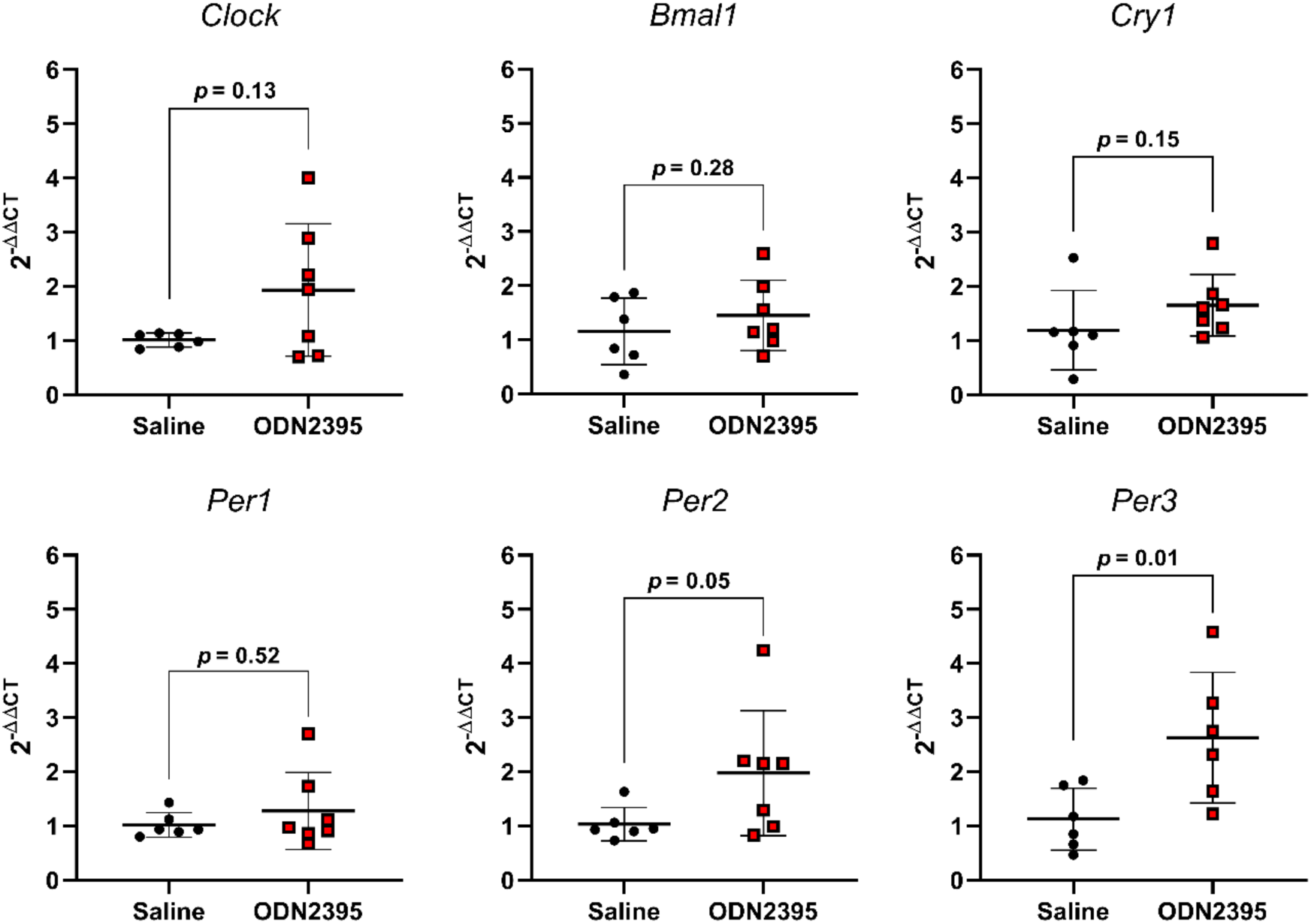
Effect of a single exposure to ODN2395 on clock gene expression in rat placentas on gestational day 14. Placental gene expression of circadian rhythm regulators circadian locomoter output cycles protein kaput (*Clock*), Brain and Muscle ARNT [Aryl hydrocarbon receptor nuclear translocator-like protein]-like protein-1 (*Bmal1*), Cryptochrome circadian regulator 1 (*Cry1*), and Period circadian protein homologs 1, 2 and 3 (*Per1*, *Per2*, *Per3*) four hours after administration of ODN2395 or saline vehicle. Values are presented as mean ± SD (n = 6-7/group) and were analyzed by unpaired t-test.

**Figure 8.**
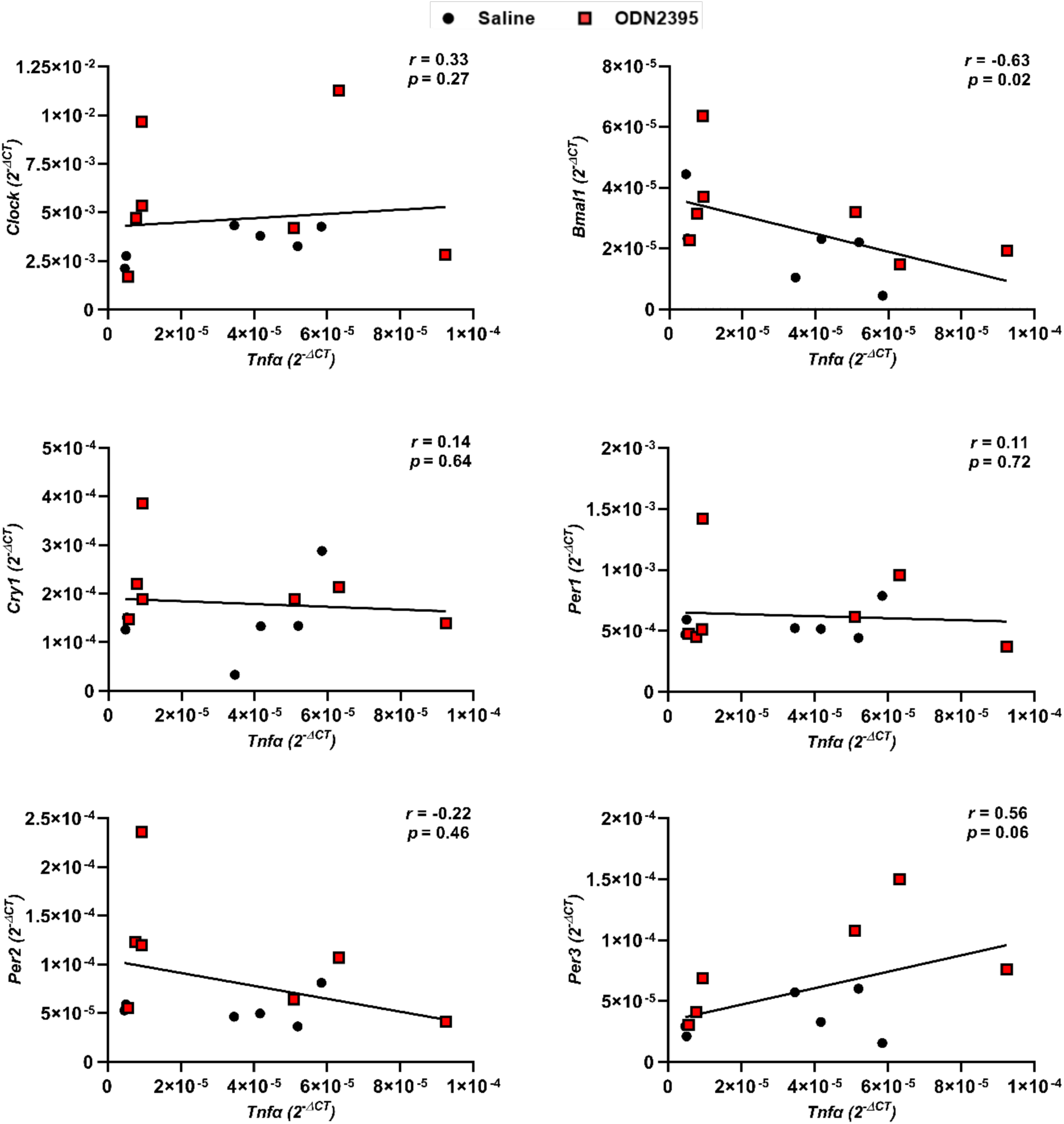
Relationship between placental proinflammatory and clock gene expression in rats administered ODN2395 or saline vehicle on gestational day 14. Spearman correlations of placental proinflammatory cytokine, Tumor necrosis factor alpha (*Tnfa*), gene expression correlated to placental clock (*Clock*, *Bmal1*, *Cry1*, *Per1*, *Per2*, *Per3*) gene expression four hours after administration of ODN2395 or saline vehicle on gestational day 14. Data normalized to *Ppia* reference gene were used for correlations (2^-ΔCT^, n = 6-7/group). Black circles = saline-administered pregnant rats, red squares = ODN2395-administered rats. CLOCK = circadian locomoter output cycles protein kaput, Bmal1 *=* Brain and Muscle ARNT-like protein, Cry1= Cryptochrome circadian regulator 1; Per1, Per2, Per3 = Period circadian protein homologs 1, 2 and 3.

## DISCUSSION

In this study, we determined the impact of exposure to CpG ODN during pregnancy on maternal circulating immune profiles, blood pressure circadian patterns, placental expression of clock network genes, and their interactive effects on perinatal outcomes in rats. We found that gestational exposure to CpG ODN disrupts maternal circadian rhythms of blood pressure and transiently affects maternal blood pressure. Additionally, exposure to CpG ODN during pregnancy induces a pregnancy-specific inflammatory profile and dysregulates expression of placental molecular clock gene expression. This dysregulation was associated with placental inflammation and might contribute to adverse effects on fetoplacental growth dynamics.

Exposure to CpG ODN during pregnancy disrupted systolic and diastolic blood pressure circadian rhythmicity. Notably, disruptions of circadian rhythms of blood pressure with or without hypertension is associated with elevated risk of cardiovascular disease (25). Indeed, human studies indicate that cardiovascular events are associated with disruptions in blood pressure circadian patterns, even in clinically normotensive individuals (26, 47, 48). Moreover, circadian rhythm disruptions of blood pressure during pregnancy are associated with pregnancy complications, such as preeclampsia, and long-term health consequences for mother and child (1, 49).

Even though circadian rhythms of blood pressure were disrupted by CpG ODN treatment, CpG ODN exposure had only transient effects on average maternal blood pressure during sleep and wake phases during the treatment window. We observed a decrease in maternal systolic blood pressure following the first treatment of CpG ODN; however, this effect was transient, which may indicate the efficiency of blood pressure regulatory mechanisms. We previously reported an increase in maternal systolic blood pressure in CpG ODN-treated pregnant rats compared to controls using the tail cuff method to obtain a single measurement on GD19, one day after the last CpG ODN injection (37, 38). The reproducibility of these results using tail cuff methodology were subsequently confirmed by others (39). The main difference between these previous studies and the current investigation is the method used for measuring maternal blood pressure. Since there are substantial differences in tail-cuff and telemetry methods (50), the discrepancies between previously published work and our current findings are not surprising. Particularly, the tail-cuff method requires restraint of the animal, which induces stress-associated responses that may contribute to stress-mediated elevations in blood pressure (51). It is possible that CpG ODN exposure renders the maternal cardiovascular system vulnerable to a “second hit”, such as prenatal stress, which is associated with pregnancy complications and adverse fetal outcomes (52). Although in the present study we did not observe an increase in maternal blood pressure, we noted that the reduction in maternal blood pressure with advancing gestational age recorded in saline-treated rats did not occur in ODN2395-treated rats. A blunted fall in maternal blood pressure has been previously observed in response to gestational stressors such as hypoxia (53, 54), high altitude (55), and exposure to inflammatory triggers (56) in various species. Thus, our data indicate that exposure to CpG ODN is a prenatal stressor that also prevents the normal reduction in blood pressure seen at the end of rodent pregnancy.

Daily (circadian) rhythms are natural internal responses finely controlled by a network of “clock” genes that respond to predictable environmental cues such as light and dark cycles that align with daily tasks, namely, physical activity and sleeping. Timing of the circadian rhythm is driven by the suprachiasmatic nucleus of the hypothalamus, which is a master central clock located within the brain that influences peripheral tissue clocks located in many organs, including the placenta, via hormonal or neuronal signals (35, 57, 58). The key molecular machinery of the circadian clock is comprised of a network of transcription factors termed “clock” genes that function in autoregulatory transcription-translation feedback loops to drive gene expression of target genes in a tissue-specific manner (59). Specifically, the transcriptional activators, Circadian locomotor output cycles kaput (CLOCK) and Brain and muscle Arnt-like protein 1 (BMAL1), induce expression of *Period* (*Per1*, *Per2*, *Per3*) and *Cryptochrome* (*Cry1* and *Cry2*) genes that act as transcriptional repressors of the clock by opposing actions of BMAL1 and CLOCK (59). In this study, we observed that a single exposure to CpG ODN at mid-pregnancy increased placental expression of clock transcriptional repressors *Per2* and *Per3*, suggesting CpG ODN influences the molecular clock network of the placental tissue.

There is growing evidence in the reciprocal relationship between clock and inflammatory gene expression and function (34, 60, 61). Thus, we assessed the impact of CpG ODN exposure on placental inflammatory gene expression. We found that CpG ODN exposure induced a pro-inflammatory response characterized by elevated placental *Tnfα* gene expression. This finding agrees with previous studies that demonstrate CpG ODN exposure during pregnancy increased placental TNF-α levels, and CpG ODN-mediated fetal demise could be rescued with blockage of TNF-α (62, 63). We then determined if CpG ODN-mediated increases in placental *Tnfα* were correlated with clock gene expression. Our data reveal increases in placental *Tnfα* are positively correlated with *Per3* expression and negatively correlated with *Bmal1* expression, suggesting a relationship between CpG ODN-mediated inflammation and clock gene expression within the placenta. Indeed, direct relationships between clock gene expression and inflammation have been identified in other tissues (61, 64). Particularly, clock genes have been shown to be negatively transcriptionally regulated by TNF-α through binding to transcriptional direct response elements (34). Our observation of negative correlations of placental *Bmal1* and *Tnfα* gene expression provides evidence that CpG ODN-mediated placental inflammation may be driving disruptions in placental clock gene expression and circadian rhythmicity. However, due to the reciprocal relationship between inflammation and clock gene expression, it is also plausible that CpG ODN-mediated disruptions in clock gene expression could be driving enhanced placental inflammation and expression of TNF-α.

Notably, placental inflammation and dysfunction are central to maternal cardiovascular disease in preeclampsia, a hypertensive disorder of pregnancy associated with disruptions in circadian rhythmicity of blood pressure and dysregulation of placental clock gene expression (65). Moreover, blood pressure is regulated by many factors produced and released by the placenta, such as hormones and vasoactive factors whose expression follows circadian patterns (35, 66). Thus, we suggest that the placenta may function as a peripheral clock that contributes to the maternal circadian rhythm of blood pressure.

Contrary to placental inflammation, our data demonstrates both a single dose of CpG ODN at the beginning of the rat third trimester and repetitive doses throughout the third trimester dampen pro-inflammatory responses in the circulation, and these immunomodulatory effects are pregnancy-specific given there were no group differences in non-pregnant animals. Moreover, a single dose of CpG ODN had greater effects on dampening proinflammatory and anti-inflammatory cytokine and chemokine responses compared to repetitive doses. This finding suggests repetitive CpG ODN exposure during pregnancy, such as occurs during rapid fetal growth and cell turnover in the third trimester, may elicit tolerance to TLR9 stimulation during late gestation. In fact, repetitive TLR stimulation has been shown to induce tolerance to stimuli (67). Even so, alterations in expression and activity of TLR9 have been demonstrated in placentas and circulation of pregnant women diagnosed with pregnancy complications such as preeclampsia (11, 68–70), and this TLR9 dysregulation may alter the impact of CpG ODN exposure on immune modulation during pregnancy. Importantly, we and others have previously demonstrated the pregnancy-specific effects of CpG ODN exposure on perinatal outcomes (37, 39). Overall, differential immunomodulatory effects of CpG ODN exposure based on reproductive status (non-pregnant vs pregnant), tissue (placenta vs circulation), or disease status (healthy or preeclampsia) may contribute to divergent responses to CpG ODN exposure, such as enhanced susceptibility to infection and reduced safety and efficacy of CpG ODN-containing vaccines administered to pregnant women. Furthermore, our findings reveal that circulating inflammatory status did not reflect placental inflammatory status, highlighting the limitation of plasma sampling of inflammatory cytokines as a proxy of placental health status.

Similar to our previous studies, there were no group differences in mean absolute placental and fetal biometrics (37, 38). Others have shown that exposure to CpG ODN during the first trimester of pregnancy in mice is teratogenic, resulting in more than 50% resorptions, fetal craniofacial malformations, and polydactyly (62, 71). These studies, however, exposed mice to much greater doses of CpG ODN (300-400 µg/dam, ∼10-20 mg/kg) and started the treatments prior to placentation. Additionally, CpG ODN exposure in the second trimester of mouse pregnancy resulted in increased rate of spontaneous abortions and fetal growth restriction (39). Previous toxicology studies suggest that litter size and resorption numbers should be taken into consideration when fetal growth is assessed in responses to a chemical substance. Our multivariate analyses revealed CpG ODN reduced fetal weights in remaining viable pups when resorptions were increased within a litter, suggesting that CpG ODN exposure impairs placental function and disrupts fetoplacental growth dynamics. Given the CpG ODN exposure occurred late in pregnancy after placentation, it is likely that CpG ODN exposure in our study impacted placental function rather than inducing anatomical defects.

### Advances, limitations, and future studies

In conclusion, our novel findings elucidate the adverse impact of unmethylated CpG DNA on maternal circadian rhythms of blood pressure, placental clock gene expression, and fetal growth. Gestational exposure to CpG DNA, such as exposure to a bacterial or viral infections, tissue turnover from rapid fetoplacental growth, or vaccination using CpG ODN adjuvants, may contribute to adverse perinatal outcomes including maternal cardiovascular complications and aberrant fetal growth and heighted maternal and offspring cardiovascular risk (Figure 9).

**Figure 9.**
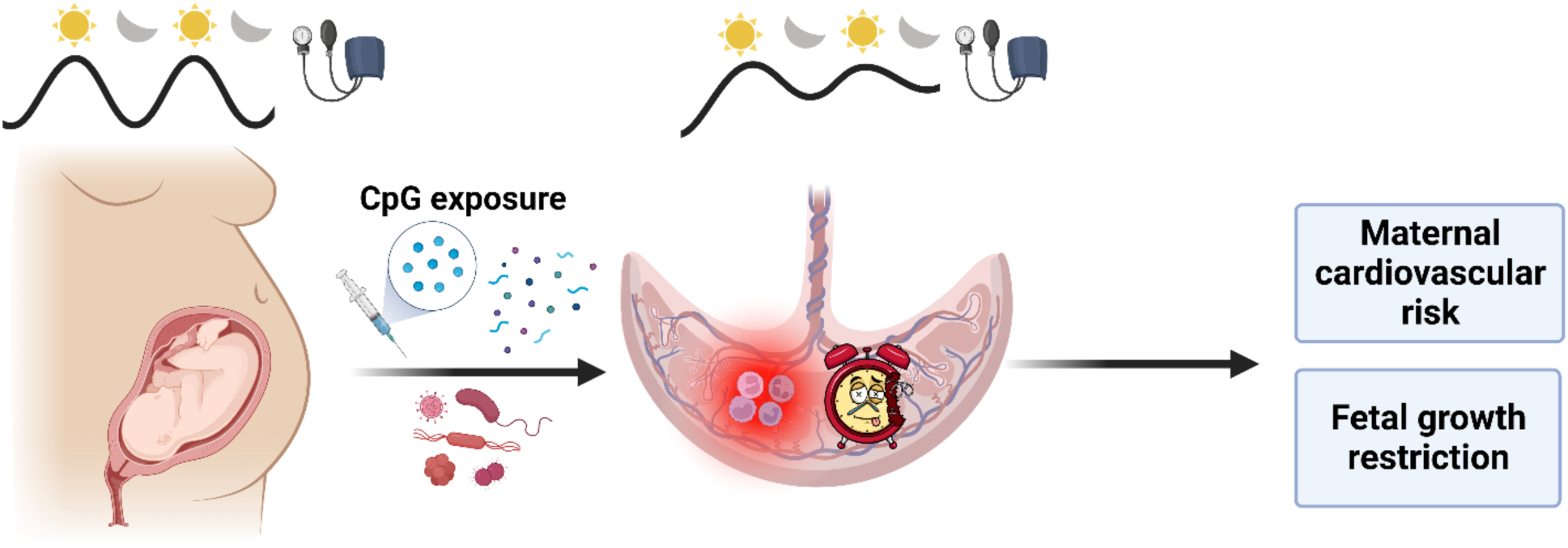
Graphical abstract. Gestational exposure to unmethylated CpG oligonucleotides dysregulates placental molecular clock network and fetoplacental growth dynamics and disrupts maternal blood pressure circadian rhythms in rats. These unmethylated CpG DNA-driven outcomes may contribute to maternal cardiovascular risk and fetal growth restriction.

Here, we characterized the placental clock gene network in response to a gestational stressor and reported an association between inflammatory and clock gene expression. These data set the foundation for future investigations to determine how immune system stimulation during pregnancy may affect circadian patterns in placental secretory function, which was not assessed in the current investigation and could have a significant impact on both the mother and the developing fetus. Additionally, our data demonstrated maternal blood pressure circadian rhythms were disrupted in response to immune system stimulation. This finding in novel and lend a framework to future studies to determine the role and contribution of central vs. peripheral oscillators to maternal blood pressure regulation during pregnancy. It is noteworthy that in this study, we stimulated the immune system pharmacologically during an otherwise healthy rat pregnancy. Future studies could interrogate the association of the maternal immune system and circadian rhythms to discover potential targets for intervention in experimental models of pregnancy complications, such as preeclampsia.

## AUTHOR CONTRIBUTIONS

JLB, SCC, and SG conceived and designed experiments. JLB, SCC, CAR, SMT, JJG, JTL, and OO conducted experiments. JLB, CAR, and SG analyzed and interpreted the data. JLB, CAR, and SG prepared figures. JLB and SG drafted the manuscript. JLB, SCC, CAR, SMT, JJG, JTL, OO, and SG edited and revised the manuscript and approved of the final version.

## ACKNOWLEDGEMENTS

Present address: Contessa A. Ricci, Washington State University, Pullman, Department of Animal Sciences, Pullman, WA, USA.

