## Supplemental Tables and Figure for "Gestational exposure to unmethylated CpG oligonucleotides dysregulates placental molecular clock network and fetoplacental growth dynamics, and disrupts maternal blood pressure circadian rhythms in rats"

**Table S1. Primer sequences for quantitative real-time PCR analysis of placental cytokine and clock gene expression.**

| Primer | Sequence (5'-3') | | Ref |
|---|---|---|---|
| *Tnfa* | *Forward:* | ACTGAACTTCGGGGTGATTG | (1) |
| | *Reverse:* | GCTTGGTGGTTTGCTACGAC | |
| *Il6* | *Forward:* | TGATGGATGCTTCCAAACTG | (1) |
| | *Reverse:* | GAGCTTGGAAGTTGGGGTA | |
| *Il1β* | *Forward:* | CACCTTCTTTTCCTTCATCTTTG | (1) |
| | *Reverse:* | GTCGTTGCTTGTCTCTCCTTGTA | |
| *Clock* | *Forward:* | ACAGCGCACACACAGGCCTTC | (2) |
| | *Reverse:* | TGGCGGCGCCCTGTGATCTA | |
| *BmalI* | *Forward:* | ACACTGCACCTCGGGAGCGA | (2) |
| | *Reverse:* | CGCCGAGCTCCAGAGCACAA | |
| *Cry1* | *Forward:* | AGCTGGCCACTGAGGCTGGT | (2) |
| | *Reverse:* | TGCTGGCATCTCCAGGGGCT | |
| *Per1* | *Forward:* | CGCACTTCGGGAGCTCAAACTTC | (2) |
| | *Reverse:* | GTCCATGGCACAGGGCTCACC | |
| *Per2* | *Forward:* | TGAGCTCCTTGGCGTTGCCG | (2) |
| | *Reverse:* | ACTCAGGCCCACTGGCCACA | |
| *Per3* | *Forward:* | TTTTCCCCTTCAAGACATGG | (2) |
| | *Reverse:* | GTCCATGGCACAGGGCTCACC | |
| *Sdha** | *Forward:* | TGGGGCGACTCGTGGCTTTC | (2) |
| | *Reverse:* | CCCCGCCTGCACCTACAACC | |

| | | |
|---|---|---|
| *Ppia** | *Forward:* AGCATACAGGTCCTGGCATC<br>*Reverse:* TTCACCTTCCCAAAGACCAC | (2) |

*, housekeeping genes included in each qPCR plate layout. *Ppia* was found to be more stable in all samples and was used for determination of comparative gene expression.

**Table S2. Determination of model with best fit for evaluating average fetal weight**

| **Model parameters** | **df** | **AIC** |
|---|---|---|
| Treatment, litter size, average placental weight | 9 | -20.08382 |
| Treatment, average placental weight, number of resorptions | 9 | **-50.29942** |
| Treatment, average placental weight, proportion of resorptions in litter | 10 | -35.06153 |

**AIC =** Akaike's Information Criterion. Bold text represents best fit (lowest AIC).

**Table S3. Pre-pregnancy blood pressure and heart rate averages during 12-hr sleep and wake cycles.**

| | | Saline | ODN2395 | *P*-value |
|---|---|---|---|---|
| **SBP** | **Sleep** | 132.9 ± 10.77 | 126.4 ± 5.09 | 0.21 |
| | **Wake** | 135.2 ± 10.03 | 126.9 ± 4.47 | 0.09 |
| **DBP** | **Sleep** | 98.95 ± 3.83 | 96.99 ± 6.50 | 0.52 |
| | **Wake** | 100.9 ± 4.45 | 97.48 ± 8.34 | 0.95 |
| **MAP** | **Sleep** | 115.8 ± 5.81 | 110.9 ± 4.87 | 0.11 |
| | **Wake** | 114.5 ± 10.13 | 111.4 ± 5.73 | 0.49 |
| **HR** | **Sleep** | 212.5 ± 11.85 | 208.4 ± 5.33 | 0.54 |
| | **Wake** | 210.0 ± 5.03 | 208.1 ± 17.64 | 0.78 |

Values presented as mean ± SD and analyzed by unpaired t-test or Mann-Whitney test (n = 6-8/group). Each 12-hour sleep or wake cycle was averaged over three days prior to mating to obtain pre-pregnancy baseline measurements. SBP = systolic blood pressure, DBP = diastolic blood pressure, MAP = mean arterial pressure, HR = heart rate.

**Table S4. Weighted multiple linear regression model assessing effects of covariates on fetal weight on gestational day 20.**

| Variable | Estimate | SE | *t*-value | *p*-value |
|---|---|---|---|---|
| (Intercept) | 1.32 | 0.63 | 2.09 | 0.09 |
| Treatment | 0.93 | 0.64 | 1.45 | 0.21 |
| Placental weight (g) | 2.08 | 1.19 | 1.74 | 0.14 |
| Resorptions | 0.42 | 0.20 | 2.09 | 0.09 |
| Treatment x Placental weight (g) | -1.87 | 1.21 | -1.54 | 0.18 |
| **Treatment x Resorptions** | **-0.86** | **0.25** | **-3.48** | **0.02** |
| Placental weight (g) x Resorptions | -0.89 | 0.38 | -2.35 | 0.07 |
| **Treatment x Placental weight (g) x Resorptions** | **1.76** | **0.51** | **3.42** | **0.02** |

SE = standard error. Multiple $R^2$: 0.9598; Adjusted $R^2$: 0.9036; model p-value: 0.003; bolded text indicates statistical significance.

**Table S5. Cytokine and chemokine protein expression across various reproductive status.**

| Cytokine/ Chemokine | Cytokine/Chemokine Function | Pregnancy Status | Treatment | Mean ± SE | p-value |
|---|---|---|---|---|---|
| TNF-α | Proinflammatory | NP | Saline | 194.3 ± 25.24 | 0.78 |
| | | | ODN2395 | 181.4 ± 35.5 | |
| | | GD14 | Saline | 139.1 ± 19.77 | ***0.02*** |
| | | | ODN2395 | 75.16 ± 11.83 | |
| | | GD20 | Saline | 349.3 ± 38.58 | 0.12 |
| | | | ODN2395 | 267.4 ± 27.15 | |
| IL-6 | Proinflammatory | NP | Saline | 20.91 ± 11.74 | 0.53 |
| | | | ODN2395 | 13.93 ± 10.79 | |
| | | GD14 | Saline | 3.14 ± 0.0 | 1.0 |
| | | | ODN2395 | 3.14 ± 0.0 | |
| | | GD20 | Saline | 310.2 ± 47.93 | 0.36 |
| | | | ODN2395 | 188.2 ± 56.47 | |
| IL-1β | Proinflammatory | NP | Saline | 13.31 ± 10.40 | 0.92 |
| | | | ODN2395 | 9.23 ± 5.361 | |
| | | GD14 | Saline | 61.28 ± 23.16 | ***0.02*** |
| | | | ODN2395 | 1.463 ± 0.0 | |
| | | GD20 | Saline | 8.56 ± 7.10 | 0.37 |
| | | | ODN2395 | 1.46 ± 0.0 | |
| IL-1α | Proinflammatory | NP | Saline | 53.38 ± 18.22 | 0.59 |
| | | | ODN2395 | 69.67 ± 23.38 | |

| | | | | | |
|---|---|---|---|---|---|
| | | GD14 | Saline | 47.25 ± 20.73 | ***0.03*** |
| | | | ODN2395 | 1.89 ± 0.0 | |
| | | GD20 | Saline | 108.2 ± 14.87 | ***0.03*** |
| | | | ODN2395 | 63.22 ± 10.38 | |
| IL-18 | Proinflammatory | NP | Saline | 428.6 ± 84.88 | 0.10 |
| | | | ODN2395 | 271.6 ± 26.66 | |
| | | GD14 | Saline | 445.1 ± 136.6 | 0.60 |
| | | | ODN2395 | 352.2 ± 104.2 | |
| | | GD20 | Saline | 2220.0 ± 348.1 | 0.40 |
| | | | ODN2395 | 1716.0 ± 476.6 | |
| IL-12p70 | Proinflammatory | NP | Saline | 74.58 ± 12.25 | 0.91 |
| | | | ODN2395 | 76.92 ± 15.74 | |
| | | GD14 | Saline | 53.33 ± 8.95 | ***0.03*** |
| | | | ODN2395 | 30.15 ± 3.70 | |
| | | GD20 | Saline | 148.2 ± 32.91 | 0.06 |
| | | | ODN2395 | 63.44 ± 21.70 | |
| IL-17A | Proinflammatory | NP | Saline | 29.18 ± 8.34 | 0.35 |
| | | | ODN2395 | 19.70 ± 2.82 | |
| | | GD14 | Saline | 21.55 ± 3.38 | ***0.01*** |
| | | | ODN2395 | 10.64 ± 1.29 | |
| | | GD20 | Saline | 25.47 ± 4.18 | ***0.03*** |
| | | | ODN2395 | 10.95 ± 4.11 | |
| IFNγ | Proinflammatory | NP | Saline | 165.1 ± 40.04 | 0.50 |

| | | | | | |
|---|---|---|---|---|---|
| | | | ODN2395 | 124.9 ± 42.30 | |
| | | GD14 | Saline | 12.66 ± 7.14 | 0.47 |
| | | | ODN2395 | 6.32 ± 4.34 | |
| | | GD20 | Saline | 174.8 ± 47.64 | 0.15 |
| | | | ODN2395 | 87.12 ± 27.75 | |
| IL-7 | Proinflammatory | NP | Saline | 19.69 ± 3.78 | 0.98 |
| | | | ODN2395 | 19.56 ± 3.21 | |
| | | GD14 | Saline | 23.18 ± 4.84 | ***0.03*** |
| | | | ODN2395 | 10.46 ± 1.84 | |
| | | GD20 | Saline | 27.36 ± 5.54 | 0.12 |
| | | | ODN2395 | 17.19 ± 5.52 | |
| IL-4 | Anti-inflammatory | NP | Saline | 141.9 ± 17.77 | 0.58 |
| | | | ODN2395 | 124.3 ± 24.10 | |
| | | GD14 | Saline | 84.85 ± 13.94 | 0.08 |
| | | | ODN2395 | 59.22 ± 5.85 | |
| | | GD20 | Saline | 138.0 ± 25.44 | 0.20 |
| | | | ODN2395 | 97.51 ± 29.12 | |
| IL-5 | Anti-inflammatory | NP | Saline | 410.5 ± 32.85 | 0.37 |
| | | | ODN2395 | 360.6 ± 41.64 | |
| | | GD14 | Saline | 309.0 ± 24.22 | ***0.04*** |
| | | | ODN2395 | 251.7 ± 12.07 | |
| | | GD20 | Saline | 496.7 ± 46.34 | 0.38 |
| | | | ODN2395 | 414.8 ± 82.07 | |

| | | | | | |
|---|---|---|---|---|---|
| IL-13 | Anti-inflammatory | NP | Saline | 102.2 ± 43.33 | 0.51 |
| | | | ODN2395 | 57.74 ± 25.69 | |
| | | GD14 | Saline | 28.76 ± 20.08 | 0.15 |
| | | | ODN2395 | 1.14 ± 0.00 | |
| | | GD20 | Saline | 76.02 ± 38.72 | 0.20 |
| | | | ODN2395 | 64.43 ± 63.29 | |
| IL-10 | Anti-inflammatory | NP | Saline | 104.6 ± 15.57 | 0.97 |
| | | | ODN2395 | 106.0 ± 26.29 | |
| | | GD14 | Saline | 88.49 ± 24.24 | 0.06 |
| | | | ODN2395 | 20.87 ± 11.25 | |
| | | GD20 | Saline | 14.28 ± 6.08 | 0.84 |
| | | | ODN2395 | 25.67 ± 14.73 | |
| IL-2 | Anti-inflammatory | NP | Saline | 762.4 ± 174.5 | 0.34 |
| | | | ODN2395 | 554.8 ± 121.6 | |
| | | GD14 | Saline | 422.4 ± 108.7 | ***0.01*** |
| | | | ODN2395 | 85.22 ± 49.52 | |
| | | GD20 | Saline | 707.2 ± 168.9 | 0.47 |
| | | | ODN2395 | 683.1 ± 406.0 | |
| GRO/KC | Chemoattractant | NP | Saline | 21.88 ± 6.68 | 0.55 |
| | | | ODN2395 | 54.03 ± 16.75 | |
| | | GD14 | Saline | 20.49 ± 9.27 | 0.61 |
| | | | ODN2395 | 7.19 ± 1.96 | |
| | | GD20 | Saline | 8.72 ± 2.50 | 0.42 |

| | | | | | |
|---|---|---|---|---|---|
| | | | ODN2395 | 9.37 ± 4.09 | |
| MCP-1 | Chemoattractant | NP | Saline | 269.2 ± 16.04 | 0.52 |
| | | | ODN2395 | 285.8 ± 15.90 | |
| | | GD14 | Saline | 396.4 ± 37.94 | ***0.03*** |
| | | | ODN2395 | 286.0 ± 21.42 | |
| | | GD20 | Saline | 150.0 ± 8.74 | ***0.01*** |
| | | | ODN2395 | 93.74 ± 17.41 | |
| MIP-1α | Chemoattractant | NP | Saline | 9.58 ± 1.78 | 0.63 |
| | | | ODN2395 | 10.62 ± 1.16 | |
| | | GD14 | Saline | 14.53 ± 2.06 | 0.71 |
| | | | ODN2395 | 13.12 ± 3.04 | |
| | | GD20 | Saline | 9.93 ± 3.21 | 0.39 |
| | | | ODN2395 | 6.62 ± 2.98 | |
| MIP-3α | Chemoattractant | NP | Saline | 13.28 ± 0.98 | 0.62 |
| | | | ODN2395 | 14.25 ± 1.57 | |
| | | GD14 | ODN2395 | 18.98 ± 2.09 | ***0.03*** |
| | | | Saline | 12.53 ± 1.45 | |
| | | GD20 | ODN2395 | 17.81 ± 1.91 | ***0.05*** |
| | | | Saline | 12.82 ± 1.18 | |
| RANTES | Chemoattractant | NP | Saline | 116.6 ± 17.90 | 0.54 |
| | | | ODN2395 | 132.0 ± 16.38 | |
| | | GD14 | Saline | 265.4 ± 35.23 | ***0.01*** |
| | | | ODN2395 | 135.0 ± 23.63 | |

| | | | | | |
|---|---|---|---|---|---|
| | | GD20 | Saline | 250.3 ± 31.66 | 0.40 |
| | | | ODN2395 | 196.1 ± 55.76 | |

Values presented as mean ± SE and analyzed by unpaired t-tests (n = 6-9/group). Data that were not normally distributed were log transformed prior to statistical comparisons. Significant *p*-values are bolded.

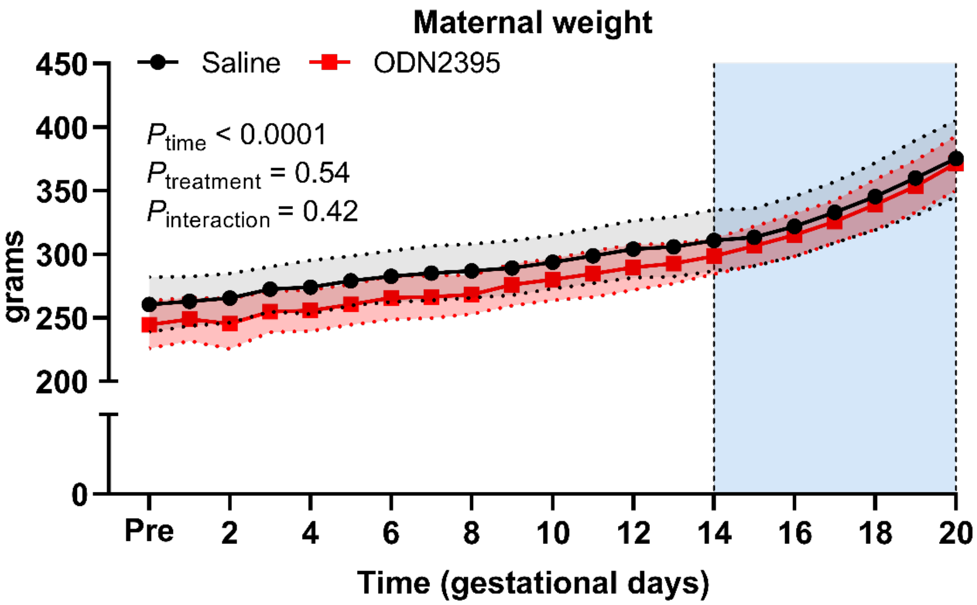


**Figure S1. Maternal weight gain throughout gestation.** ODN2395 administration had no effect on maternal body weight gain during pregnancy. Analyzed using repeated measures Two-Way ANOVA with Sidak's multiple comparisons post-hoc analysis. All values are presented as mean ± SD. N = 7-9/group. PRE, average of 3 days prior to mating. Shaded vertical region represents treatment window.
